# Comparison of human population receptive field estimates between scanners and the effect of temporal filtering

**DOI:** 10.1101/696690

**Authors:** Catherine Morgan, D Samuel Schwarzkopf

**Affiliations:** School of Psychology and Centre for Brain Research, The University of Auckland, New Zealand; Brain Research New Zealand; School of Optometry & Vision Science, Faculty of Medical & Health Sciences, University of Auckland, New Zealand; Experimental Psychology, University College London, United Kingdom

## Abstract

Population receptive field (pRF) analysis with functional magnetic resonance imaging (fMRI) is an increasingly popular method for mapping visual field representations and estimating the spatial selectivity of voxels in human visual cortex. However, the multitude of experimental setups and processing methods used makes comparisons of results between studies difficult. Here, we show that pRF maps acquired in the same three individuals using comparable scanning parameters on a 1.5 and a 3 Tesla scanner located in two different countries are very similar. As expected, the signal-to-noise ratio for the 3 Tesla data was superior; critically, however, estimates of pRF size and cortical magnification did not reveal any systematic differences between the sites. Moreover, we tested the effect of low-pass filtering of the time series on pRF estimates. Unsurprisingly, filtering enhanced goodness-of-fit, presumably by removing high-frequency noise. However, there was no substantial increase in the number of voxels containing meaningful retinotopic signals after low-pass filtering. Importantly, filtering also increased estimates of pRF size in the early visual areas which could substantially skew interpretations of spatial tuning properties. Our results therefore suggest that pRF estimates are generally comparable between scanners of different field strengths, but temporal filtering should be used with caution.

**Precis:** Population Receptive Field mapping performed with similar protocols at two different sites, a 1.5T MRI scanner in London, and a 3T scanner in Auckland, yielded comparable results. Temporal filtering of the fMRI time course increased concordance of modelled pRFs, but introduced a bias in pRF size.

## Introduction

Population receptive field analysis with fMRI has become a popular method in the toolbox of visual neuroscience. It has not only been used for mapping cortical organization (Dumoulin and Wandell, 2008; Amano et al., 2009; Winawer et al., 2010), but also to study spatial integration in the visual cortex (Dumoulin et al., 2014; Harvey and Dumoulin, 2016), reveal the effects of attention on visual processing (de Haas et al., 2014; Klein et al., 2014, 2018; Kay et al., 2015; Vo et al., 2017), show differences in patients and special populations (Levin et al., 2010; Hoffmann et al., 2012; Schwarzkopf et al., 2014; Clavagnier et al., 2015; Anderson et al., 2017) and for reconstructing the neural signature of perceptual processes (Kok and de Lange, 2014; Kok et al., 2016a, 2016b; Ekman et al., 2017; Senden et al., 2019). The most wide-spread technique involves fitting a two-dimensional symmetric Gaussian model of the pRF to the time series of each voxel in visual cortex responding to a set of stimuli (Dumoulin and Wandell, 2008). Estimates of pRF position and size reflect an aggregate of the position preferences and sizes of thousands of neuronal receptive fields of the cells within the imaging voxel, and also incorporates extra-classical receptive field interactions. Further, in fMRI as underlying neuronal activity is inferred through neurovascular coupling, pRF measurements are affected by hemodynamic factors (although it has been shown that pRFs estimated from fMRI data have a close correspondence with receptive field properties in electrophysiological experiments (Dumoulin and Wandell, 2008; Winawer et al., 2011; Alvarez et al., 2015; but see also Keliris et al., 2019).

The indirect nature of estimating pRFs from fMRI data suggests therefore that there could be considerable variability in derived measurements. Direct test-retest evaluations with the same experimental setup have shown that pRF mapping experiments are robust and repeatable (Senden et al., 2014; van Dijk et al., 2016; Benson et al., 2018). However, for *different* experimental setups, for instance, in terms of the magnetic field strength and the particular pulse sequence used to acquire fMRI data the comparability has not been assessed. The signal-to-noise ratio of MRI is proportional to voxel volume and the strength of the static magnetic field (Edelstein et al., 1986), and hence pRF measurements at higher magnetic field strength might be more accurate. However, the temporal resolution (or repetition time, TR, of image acquisition), directly affects the contribution of different noise frequencies and the contribution of physiological nuisance factors like respiration or cardiac pulsation. Noise in fMRI data therefore has multiple contributions (physiological, thermal and system related) and the relationship between the temporal signal to noise ratio (TSNR) in a fMRI time course has a non-linear relationship with static SNR (Murphy et al., 2007), i.e. gains in static SNR, from for example increased field strength, may not translate to proportional gains in TSNR due to a limit where physiological noise dominates.

Here, we conducted pRF mapping on three individuals using identical TR and voxel size on a 1.5 Tesla (1.5T) scanner in London, United Kingdom and a 3 Tesla (3T) scanner in Auckland, New Zealand. Critically, our aim was not to test the effect of magnetic field strength alone, but rather to compare pRF estimates from typical methods used at each scanning facility that also *includes* a different field strength. Thus, our findings should reflect a relatively conservative estimate of the variability of pRF parameters under comparable conditions at the two sites.

Typical pRF studies evaluate the goodness-of-fit of the pRF model by means of the coefficient of determination (R^2^) of the correlation between the observed and predicted time series. This measure however strongly depends on the temporal resolution of the signal acquisition. In our recent experiments we have used an accelerated multiband sequence with 1 s TR (Moutsiana et al., 2016, 2018; van Dijk et al., 2016; de Haas and Schwarzkopf, 2018; Hughes et al., 2019). At this temporal resolution, there is a considerable contribution of high-frequency noise to the signal (Figure 1). Since our analysis does not typically involve any temporal filtering beyond linear detrending (essentially, a wide-bandwidth high-pass filter removing only very low frequencies that are attributed to slow drifts in the signal), this probably explains why the overall R^2^ in our studies is comparably low compared to those reported by others who use more standard fMRI TRs of 2-3 s. We therefore conducted an analysis comparing pRF parameters obtained using our standard analysis (minimal filtering), to filtering with two low-pass filters that approximate longer TR acquisition.

**Figure 1.**
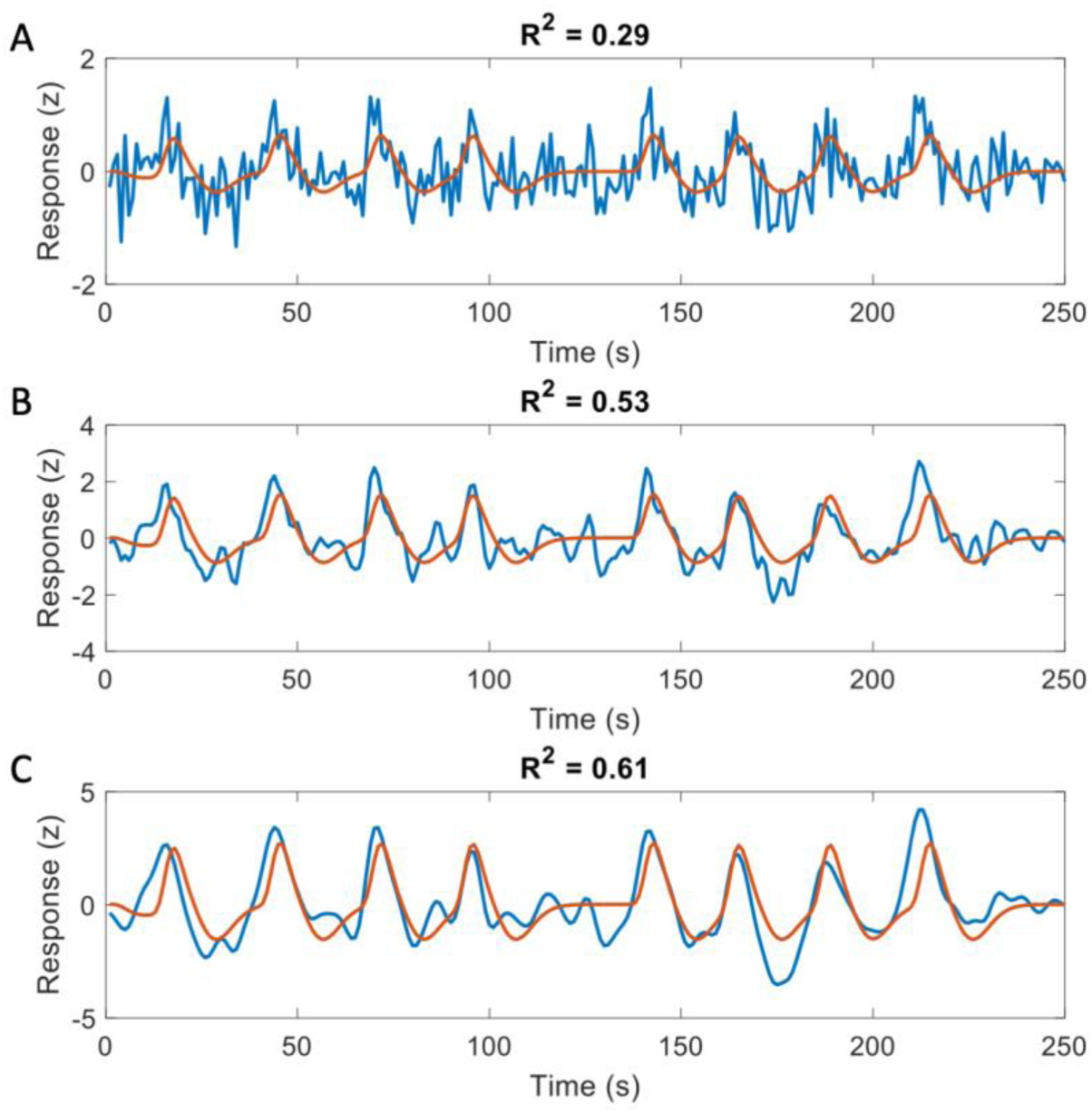
Low-pass filtering the time series improves goodness-of-fit quantified by R^2^. The same predicted time series (red curves) is shown overlaid in the unfiltered time series (A), and low-pass filtered time series with kernel 1 s (B) or 2 s (C). R^2^ is substantially larger for the filtered time series because high-frequency noise has been removed.

## Methods

### Participants

Three adult volunteers (2 female, 1 left-handed, aged 30, 39 and 47) participated in this study, henceforth referred to as P1, P2, and P3. They gave written informed consent to take part and all procedures were approved by local ethics review boards at University College London and the University of Auckland.

### Procedure

Participants were scanned twice with a pRF mapping protocol. The first scan took place inside a MAGNETOM Avanto 1.5T MRI scanner (Siemens Healthcare, Erlangen, Germany) at the Birkbeck/UCL Centre for NeuroImaging (BUCNI) in the Experimental Psychology department of University College London, United Kingdom (henceforth referred to as London site). The second scan took place several months later in a MAGNETOM Skyra 3T MRI scanner (Siemens Healthcare, Erlangen, Germany) at the Centre for Advanced Magnetic Resonance Imaging (CAMRI) in the Faculty of Medical & Health Sciences of the University of Auckland, New Zealand (henceforth referred to as Auckland site). In both scans, participants were scanned with six runs for pRF mapping lasting 4 min 20 s each during which functional echo-planar images were acquired. Moreover, at both centres a structural T1 weighted brain image was acquired although the structural image from the second scan (at 3T) was not used in any further analysis. During the scans, participants were instructed to remain as still as possible and fixate continuously on a small dot in the centre of the screen. They were instructed to press a button whenever the fixation dot changed colour.

### Stimuli

Stimuli were generated and presented using MATLAB (Mathworks) and Psychtoolbox 3 (Brainard, 1997) at a resolution of 1920 * 1080. At both sites, the screen subtended 34° by 19° of visual angle. Stimuli were presented on a screen at the back of the bore via a mirror mounted on the head coil. At the London site, stimuli were projected onto a screen in the back of the bore while in the Auckland site stimuli were presented on an MRI compatible 32′ widescreen LCD (Cambridge Research Systems Ltd).

At both sites, the stimuli generated were matched, and comprised bars containing a dynamic high-contrast ripple pattern as used in previous studies (de Haas et al., 2014; Schwarzkopf et al., 2014; Alvarez et al., 2015; Anderson et al., 2017) on a uniform grey background. Bars traversed the visual field in a regular sequence (e.g. from the bottom to the top), jumping by 0.38° every second. Each sweep of the bar lasted 25 s and so there were 25 jumps of the bar. Each run started with a sweep from the bottom to the top and then the sweep direction was rotated by 45° clockwise on the next sweep. There were thus eight sweeps covering a complete rotation. After the fourth and eighth sweep, a 25 s baseline period (no bars) was presented.

Bars were always 0.53° wide and at the longest (when crossing the centre of the visual field) subtended the full screen height, but because they were presented only within a circular region (diameter: 19°, i.e. the height of the screen) they were accordingly shorter at the start and end of each sweep. The outer edge of the ripple stimulus was smoothed by ramping it down to zero over 0.1°. Similarly, the stimulus contrast ramped down to zero from 0.53° to 0.43° eccentricity, thus creating a blank hole around fixation.

A blue dot (diameter: 0.09°) was present in the centre of the screen throughout each scanning run. The whole run was divided into 200 ms epochs. At each epoch, there was a 0.01 probability that the dot could change colour to purple with the constraint that no such colour changes could occur in a row. Participants were instructed to fixate the dot and press a button on a magnetic resonance-compatible button box whenever it changed colour. In addition, a radar screen pattern comprising low-contrast radial and concentric lines around fixation was presented at all times to aid fixation compliance (see (van Dijk et al., 2016). The fixation dot and radar screen pattern were also presented during the baseline period.

Prior to each run, we collected 10 dummy volumes to allow ample time for steady-state magnetisation to be reached. During this time, only the fixation dot was presented on a blank grey screen.

### Scanning parameters

At both sites, we used a 32-channel head coil where we removed the front elements because it impeded the view of the stimulus. This resulted in 20 effective channels covering the back and the sides of the head. Six pRF mapping runs of 260 T2 *-weighted image volumes were acquired (including the 10 dummy volumes). We used 36 transverse slices angled to be approximately parallel to the calcarine sulcus (planned using the T1-weighted anatomical image). At both sites, we used an accelerated multiband sequence (Breuer et al., 2005; Moeller et al., 2010) at 2.3 mm isotropic voxel resolution, field of view 96×96, and a TR of 1 s. At the London site, the scan had an echo time (TE) of 55 ms, flip angle of 75°, and a multiband/ slice acceleration factor of 4 and rBW 1628 Hz/pixel. At the Auckland site, the scan had a TE of 30 ms, flip angle of 62°, a multiband/slice acceleration factor of 3, an in-plane/parallel imaging acceleration factor of 2 and rBW was 1680 Hz/Px. After acquiring the functional data at the London site, the front portion of the coil was put back on to ensure maximal signal-to-noise levels for collecting a structural scan (a T1-weighted anatomical magnetization-prepared rapid acquisition with gradient echo scan with a 1 mm isotropic voxel size and full brain coverage).

### Data preprocessing

Functional data were preprocessed in SPM12 (Welcome Centre for Human NeuroImaging). The first 10 dummy volumes were removed. Then we performed mean bias intensity correction, realignment and unwarping of motion-induced distortions, and coregistration to the structural scan acquired at the London site using default parameters in SPM12. Using FreeSurfer (http://surfer.nmr.mgh.harvard.edu) we further used the structural scan for automatic segmentation and reconstruction as a three-dimensional surface mesh of the pial and grey-white matter boundaries (Dale et al., 1999; Fischl et al., 1999). The grey-white matter surface was then further inflated into a smooth model and a spherical model.

All further analysis was conducted using our custom SamSrf 6 toolbox (Schwarzkopf et al., 2018). Functional data were projected from volume space to the surface mesh by finding the nearest voxel located halfway between each vertex in the pial and grey-white matter surface mesh. The time series at each vertex was then linearly detrended and z-standardized before being averaged across the six runs at each scanning site.

For the temporal filtering analysis, we then further convolved the time series of each vertex with a Gaussian filter with standard deviation of 1 or 2 s, respectively.

### pRF modelling

We modelled pRFs as a symmetric, two-dimensional Gaussian defined by x_0_ and y_0_, the Cartesian coordinates of the pRF centre in visual space, and the standard deviation of the Gaussian, σ, as a measure of the pRF size. The pRF model predicted the neural response at each TR of the scan by calculating the overlap of the mapping stimulus with this Gaussian pRF profile. A binary mask of 100-by-100 pixels indicated where the stimulus appeared on the screen for each time point. The neural responses were then determined by multiplying each frame of the stimulus mask with the pRF profile and summing over the 10,000 pixels. Subsequently, the time series was convolved with a canonical hemodynamic response function determined from previous empirical data (de Haas et al., 2014) and z-standardized.

The pRF parameters at each vertex were fit using a two-stage procedure. First, we applied a coarse fit which involved an extensive grid search by correlating the actually observed time series against a set of 7650 predicted time series derived from a combination of x_0_, y_0_, and σ covering the plausible range for each parameter (see (Moutsiana et al., 2016; van Dijk et al., 2016). The parameters giving rise to the maximal correlation were then retained for the second stage, the fine fit, provided the squared correlation, R^2^, exceeded 0.01. The fine fit entailed an optimization procedure (Nelder and Mead, 1965; Lagarias et al., 1998) to refine the three pRF parameters to further maximize the correlation between observed and predicted time series. Subsequently, we used linear regression between the observed and predicted time series to fit the amplitude, β_1_, and the baseline intercept, β_0_. Up until this point, all analyses used raw data without any smoothing or interpolation. However, we then applied a Gaussian smoothing kernel (full width at half maximum = 3 mm) to the final pRF parameter maps on the spherical surface mesh. These smooth maps were used for visualization purposes and delineation of the visual areas. Moreover, they are necessary for calculating the local cortical magnification factor (CMF; Harvey and Dumoulin, 2011) because it requires a smooth visual field map without gaps and scatter. For this we determined the cortical neighbours of each vertex and calculated the area subtended by the polygon formed by their pRF centres. The same procedure was used to determine the cortical surface area (calculation performed by FreeSurfer). To calculate the CMF, we then divided the square root of cortical area by the square root of visual area.

In addition, for the data comparing the London and Auckland sites we also calculated the noise ceiling as an estimate of the maximum goodness-of-fit that could theoretically be achieved from the data of each voxel. For this, we split the six pRF mapping runs from each site into even and odd numbered runs and averaged them separately. We then calculated the Pearson correlation between these split time series, r_obs_’, and used the Spearman-Brown prophecy formula (Brown, 1910; Spearman, 1910) to determine, r_obs_, the expected reliability for the average of all six runs:

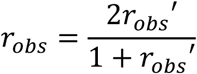

The theoretical maximum observable correlation between the predicted and observed time series, ρ_o_, for each voxel is given by the square root of this correlation (Spearman, 1904) because

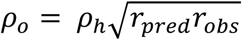

where *r*_*pred*_ is the reliability of the predictors, which is 1, and we assume the hypothetical correlation between the observed and predicted time series, ρ_h_, also assumed to be 1. The noise ceiling, the maximum observable goodness-of-fit ρ_o_^2^ is therefore equal to r_obs_.

### Further analysis

We manually delineated visual areas V1-V4 and V3A using smoothed maps from the London site by determining the borders between regions from the polar angle reversals (Sereno et al., 1995) and then also applied these delineations to the maps generated in Auckland. For each region of interest (ROIs) we then extracted pRF data and binned them into eccentricity bands 1° in width, starting from 0.5° and increasing up to 9.5°. For each bin, we calculated the mean pRF size, and the median CMF, R^2^, noise ceiling ρ_o_^2^, or the normalized goodness-of-fit, that is R^2^ divided by ρ_o_^2^. To bootstrap the dispersion of these summary statistics, we resampled the data in each bin 1000 times with replacement and determined the central 95% of this bootstrap distribution as a confidence interval.

## Results

### Comparison between scanning sites

We first compared visual field maps from the two sites, London 1.5T and Auckland 3T, by visual inspection. Figure 2 shows polar angle and eccentricity maps of the left hemisphere of one participant from both sites. It is immediately apparent that more voxels survive statistical thresholding (R^2^>0.1) for the 3T data. The cortical territory occupied by visual field maps is somewhat more extensive and more complete. Nevertheless, the orderly organization of V1-V4, V3A, as well regions in the LO complex is clearly visible in both scans, and their borders are very similar. We further quantified the map similarity by calculating correlations across all voxels above threshold in both scans (circular correlation for polar angle, Spearman’s ρ correlation for eccentricity). This showed that the polar angle maps were well correlated for two participants (P1: 0.71; P2: 0.71) although the correlation was not as strong in the final participant (P3: 0.44). Eccentricity maps were strongly correlated in all three participants (P1: 0.70; P2: 0.75; P3: 0.67).

**Figure 2.**
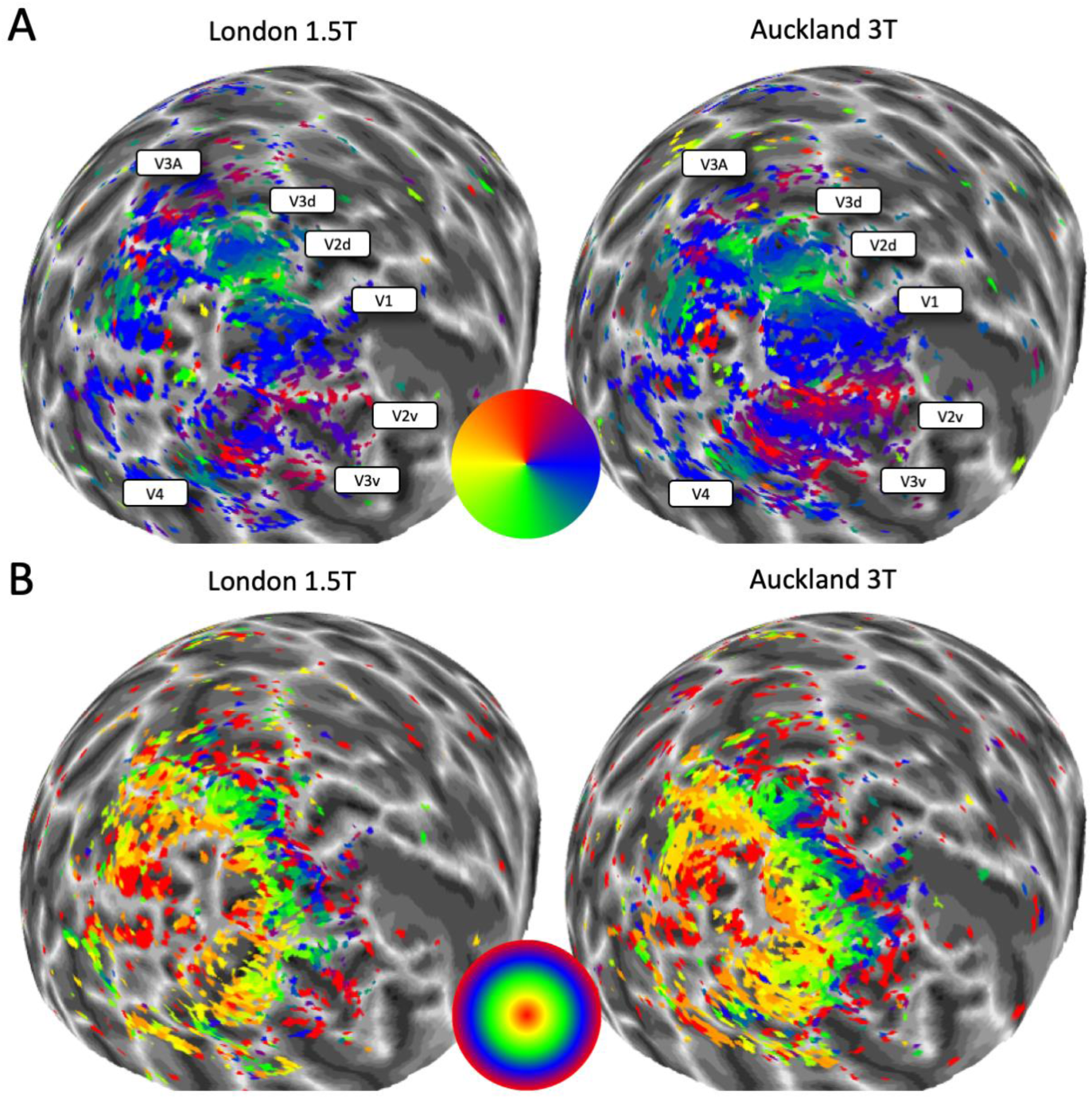
Polar angle (A) and eccentricity (B) maps of one participant from the London 1.5T site (left) and the Auckland 3T site (right) displayed on a spherical model of the left hemisphere. Colour wheels denote the pseudo-colour code for visual field maps. No smoothing or interpolation was applied to the mapping data. Voxels were thresholded at R^2^>0.1. In A, the position of the visual regions have been labelled for reference. The greyscale indicates the cortical folding pattern.

Next, we compared pRF size using 1° wide eccentricity bands to bin the data from each site and then calculating the mean for each bin. We observed that pRF size consistently increased across eccentricities and also along the visual pathway as expected. But crucially, while pRF size varied somewhat between sites, this difference was not systematic across the three participants or the five regions of interest (Figure 3A). In V4 and V3A there was somewhat greater variance at some eccentricities, likely due to the smaller size of these regions compared to V1-V3. Overall, pRF sizes were however very similar between sites. Similarly, the median cortical magnification factor (CMF) showed the expected exponential decrease from the central to the peripheral visual field. CMF curves were very comparable for the two sites (Figure 3B), except for somewhat greater CMF at 3T in V1 and V2 in the very central visual field of participants P2 and P3.

**Figure 3.**
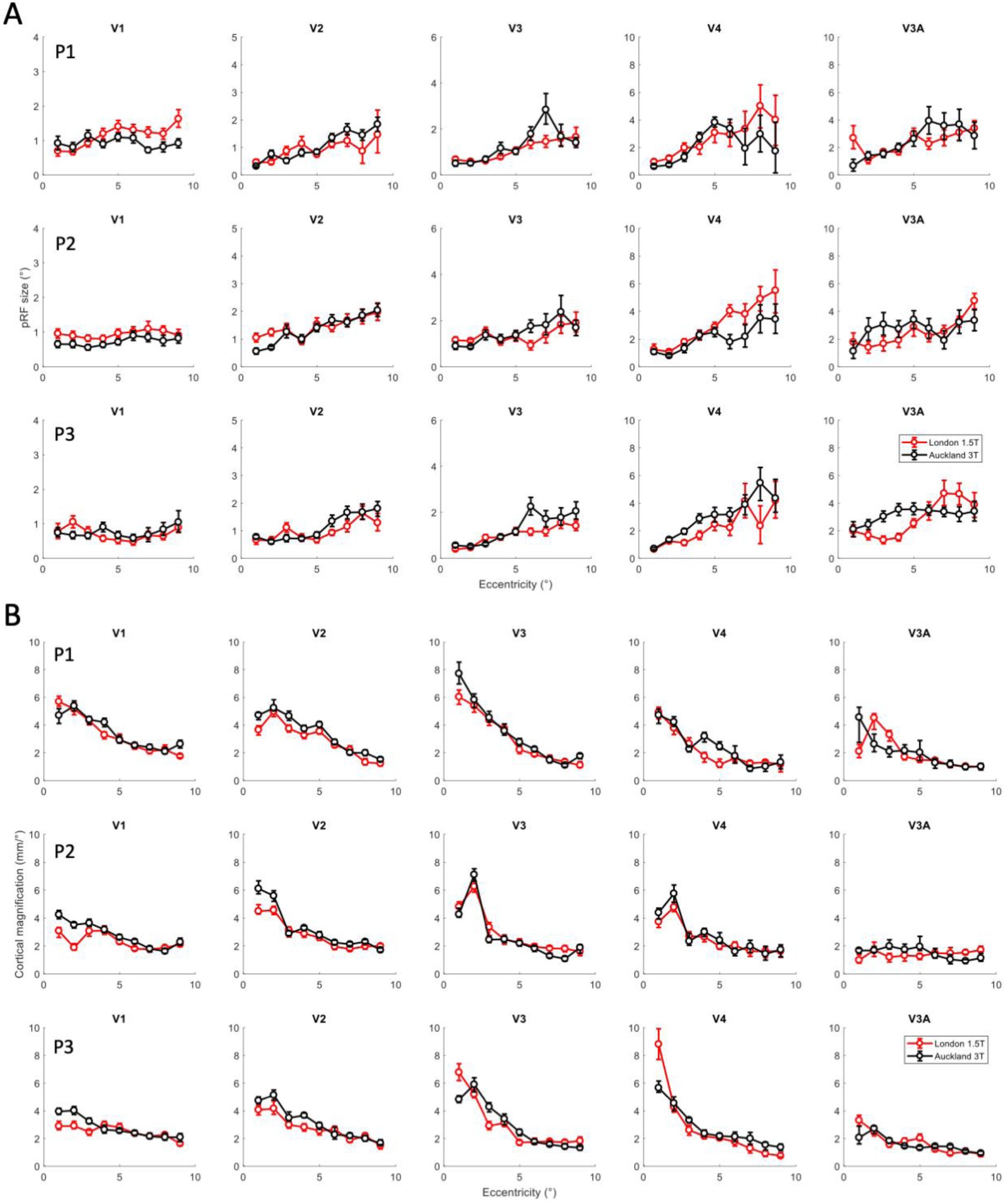
Mean pRF size (A) and median CMF (B) binned into eccentricity bands for the London 1.5T site (red) and the Auckland 3T site (black). Panels in columns show different visual regions. Panels in rows show the three participants. Error bars denote 95% confidence intervals based on bootstrapping.

We then repeated this analysis for the goodness-of-fit of the pRF model across eccentricities (Figure 4A). This showed that in V1 and V2 goodness-of-fit was notably greater for the Auckland 3T site than the London 1.5T site in all 3 participants. In higher extrastriate regions, the pattern of results was less clear, although at least for P3 goodness-of-fit was greater also in V3 and V4 (although note that in V4 for P1 very little data with very low model fits was present beyond 6° eccentricity in the Auckland data). In V3A, the model fits at both were similar, but generally lower than in the other regions and with greater variability. However, this was presumably largely driven by the overall signal-to-noise ratio. When we normalized model fits relative to the noise ceiling, ρ_o_^2^, the maximum goodness-of-fit that could theoretically be achieved given the data from each site, we found no systematic difference in goodness-of-fit between sites (Figure 3B). The curves mostly overlapped for P1 and P2 except in V1 where the London data outperformed the Auckland data. In contrast, for P3 normalized model fits for the Auckland site outperformed the London site in all regions except V3A. The noise ceiling itself was consistently higher for the Auckland data than the London data (Figure 4C).

**Figure 4.**
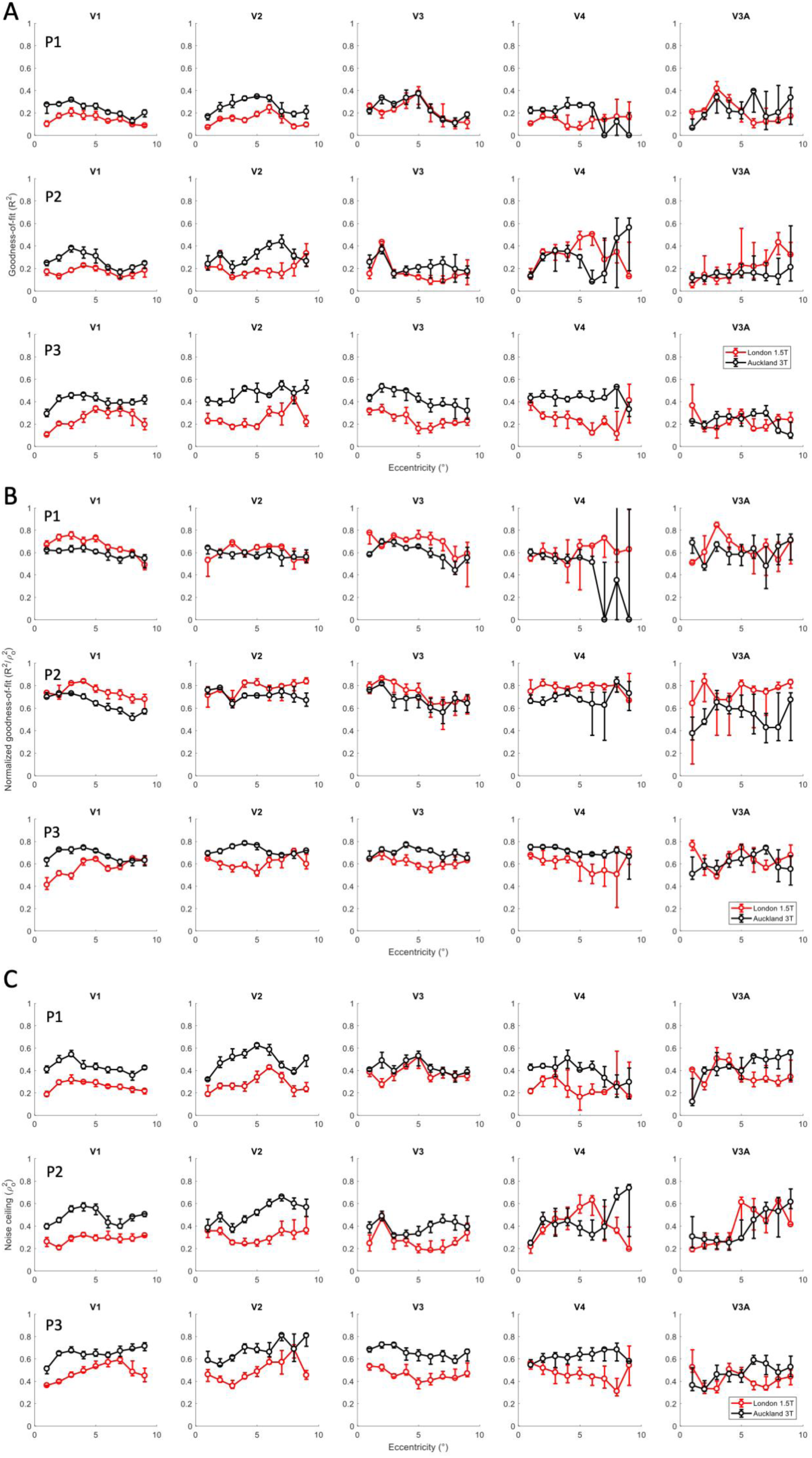
Median goodness-of-fit (A), normalized goodness-of-fit (B), and noise ceiling (C) binned into eccentricity bands for the London 1.5T site (red) and the Auckland 3T site (black). Panels in columns show different visual regions. Panels in rows show the three participants. Error bars denote 95% confidence intervals based on bootstrapping.

### Effect of temporal filtering

Goodness-of-fit of the modelled time course derived from pRF estimates and the measured fMRI time course are typically quantified by R^2^. However, the pRF model does not account for the high frequency noise observed in fast temporal resolution acquisitions like the 1 s TR we used here, which therefore likely results in lower R^2^ values. Theoretically, high frequency signals could be removed from the data and thus the model fits artifactually improved (Figure 1). We therefore reanalysed the data from the 3T site after temporal low-pass filtering the time series of each voxel by convolving it with a Gaussian kernel of either 1 s or 2 s standard deviation (e.g. Figures 1B and 1C, respectively).

Figure 5 shows visual field maps from the left hemisphere of one participant using the raw data and those after low-pass filtering. A very similar map structure is apparent in all three images. There are however also a lot of noise voxels surviving thresholding (R^2^>0.1) outside the general visual field maps. Conversely, while filtering filled in a few missing voxels within the maps, filtering did not affect the overall structure and did not increase the cortical territory of the responsive region.

**Figure 5.**
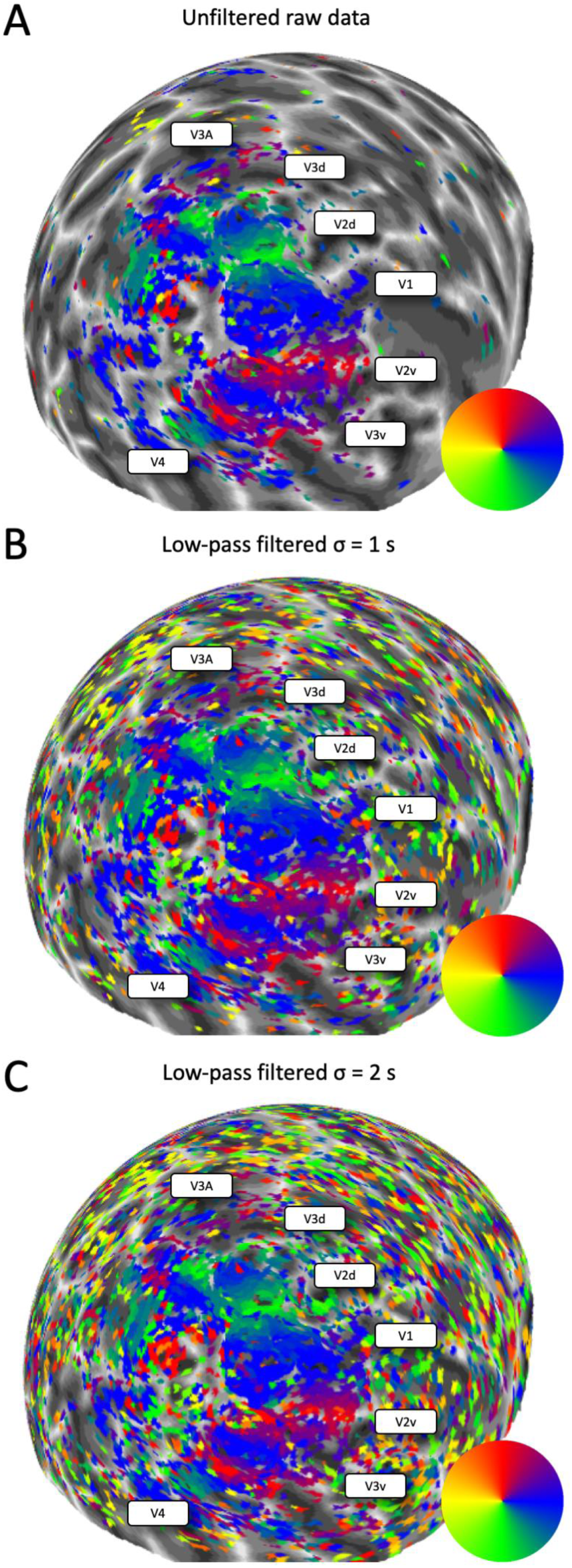
Polar angle maps of one participant from the Auckland 3T site displayed on a spherical model of the left hemisphere. Colour wheels denote the pseudo-colour code for visual field maps. No smoothing or interpolation was applied to the mapping data. Voxels were thresholded at R^2^>0.1. The position of the visual regions have been labelled for reference. The greyscale indicates the cortical folding pattern. A. Unfiltered data. B. Low-pass filtered with kernel 1 s. C. Low-pass filtered with kernel 2 s.

Next, we quantified mean pRF size and compared this for the three data sets (Figure 6A). In the early areas V1-V3, pRF size is consistently greater for the filtered time series, especially at the longer kernel of 2 s. In V4 and V3A, pRF size remained relatively similar between conditions. We also quantified the median goodness-of-fit of the pRF model (Figure 6B) and observed consistently greater model fits for both the low-pass filtered time series in V1 to V4. A similar pattern could be seen in V3A but results were generally more variable, especially in P1.

**Figure 6.**
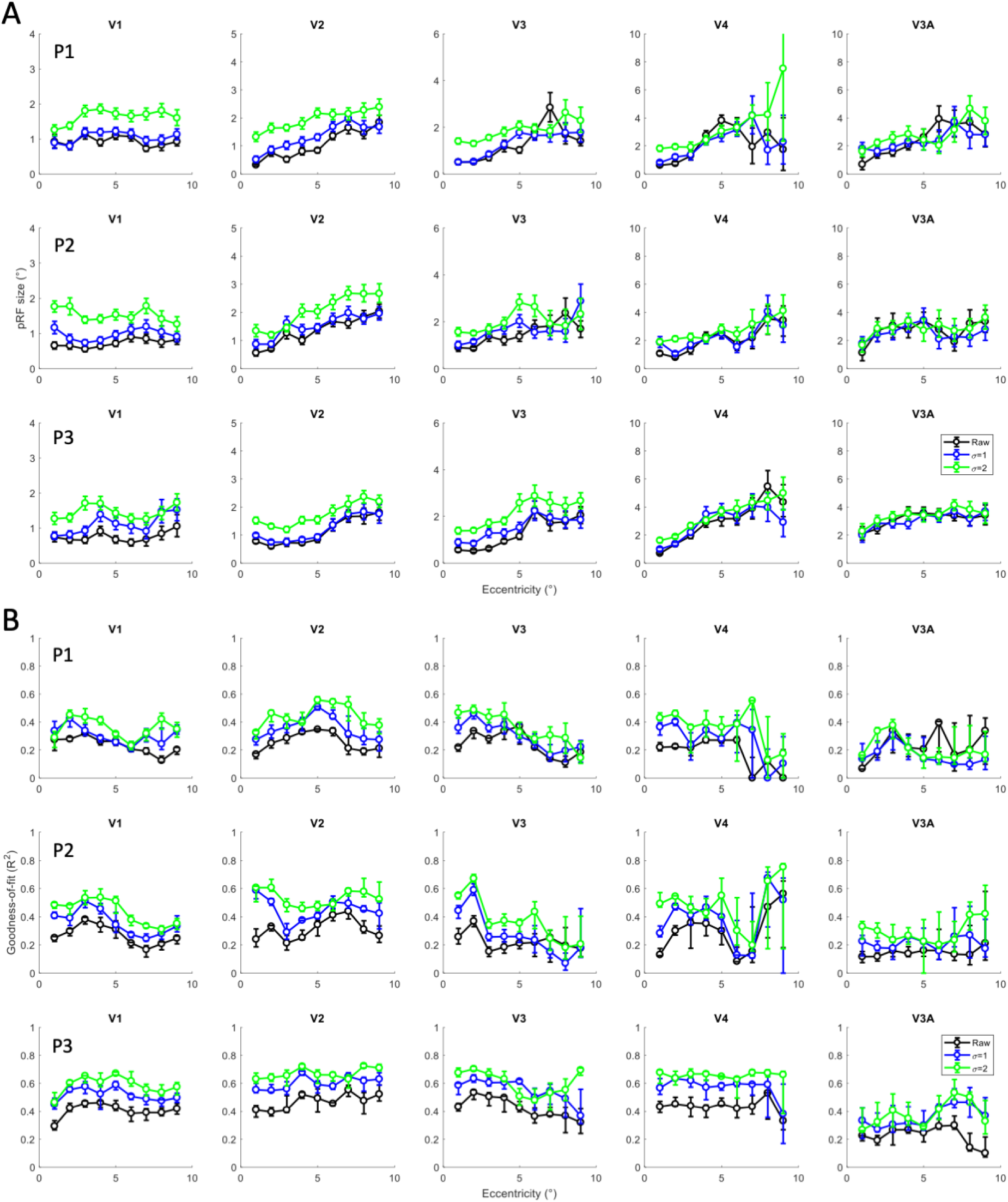
Mean pRF size (A) and median goodness-of-fit (B) binned into eccentricity bands for the Auckland data without filtering (black), and low-pass filtering with kernel 1 s (blue) and 2 s (green). Panels in columns show different visual regions. Panels in rows show the three participants. Error bars denote 95% confidence intervals based on bootstrapping.

## Discussion

Here, we compared pRF mapping data acquired using the same stimuli and participants and under comparable conditions at two sites, a 1.5T scanner in London, United Kingdom, and a 3T scanner (by the same vendor) in Auckland, New Zealand. Generally, pRF model fits were better in the Auckland data, probably in large part because of the approximately double signal-to-noise ratio from the greater static magnetic field. However, despite this difference in accuracy of data fitting, we found that visual field maps and actual estimates of pRF size and local cortical magnification were very similar between the two sites.

Although the London data were always acquired first, this is unlikely to explain our findings. All our participants were familiar with the fMRI environment, including undertaking previous pRF mapping studies. Therefore, it is unlikely that there should have been substantial training effects that could for example have changed the amount of head motion or fixation compliance between sessions. The exact parameters of the pulse sequences used and general differences in the image quality of the two scanners could of course also be contributing factors to inter-site differences. We did not apply any correction of distortions caused by the static magnetic field (Hutton et al., 2002) to either site data. Moreover, the exact slice positioning, and thus how the voxel grid resolved the grey matter, may also have differed between sites. Relevant to this, in our previous work quantifying the test-retest reliability of pRF maps we found that reliability was greater for scanning sessions in close succession on the same day than for sessions on different days (van Dijk et al., 2016). This could have been caused by fluctuations in the scanner hardware itself but also relate to differences in head position at set up. While in both cases participants were removed from the scanner between the repeat sessions, it is highly likely that positioning was more similar for the sessions conducted on the same day.

While we took painstaking steps to match the visual displays at the two sites as closely as possible, due to constraints of the experimental setup there may have been small differences between the stimuli presented. In London, images were projected onto a screen and this necessitated focussing and scaling the projected image to be of the exact size. The image in London may have been somewhat blurrier and the viewing angle more variable than in Auckland where we used a clear liquid crystal display that was always placed at the exact same position at the back of the bore. Naturally, the exact viewing angle of the stimuli also depends on the viewing distance and there may have been subtle variation between the sites although this is unlikely to have produced any systematic differences.

Nevertheless, the general extent of the visually responsive area of cortex containing clear retinotopic maps was very comparable across the sites, as were estimates of spatial tuning and cortical magnification. Therefore, we conclude that pRF estimates are very robust across scanning sites, even when using different magnetic field strengths.

At the short TR of 1 s as used in our study, noise contributes high-frequency signals to the fMRI response that are irrelevant to the sluggish blood oxygen level dependent activity the pRF model seeks to characterize. This explains why the model fits in most of our studies are relatively low compared to those reported in the literature using more conventional TRs. We therefore also sought to test what effect temporal low-pass filtering had on pRF parameter estimates. Unsurprisingly, low-pass filtering enhanced the goodness-of-fit of pRF models. However, while this filled in a few gaps in the maps it largely boosted the number of noise voxels outside the visually responsive cortex.

Crucially, low-pass filtering also generally enlarged estimates of pRF size in the early visual regions V1-V3 while data in higher regions were mostly unaffected. The pRF size estimates for the unfiltered data accord well with previous research and what one would expect from electrophysiological recordings (Dumoulin and Wandell, 2008; Harvey and Dumoulin, 2011; Winawer et al., 2011; Alvarez et al., 2015). Therefore, the pRF sizes of the filtered data presumably reflect an overestimate that seems unrealistically high (e.g. for P2 mean pRF size in central V1 was close to 2° for the most heavily filtered data).

The reason why filtering increased pRF sizes in early areas is probably the fact that filtering blurred together signals from stimuli presented close in time (proximal bar positions were presented only a few seconds apart). In the early regions where pRFs are small this would therefore be modelled as a larger pRF, whereas in higher areas in which pRFs are large enough to encompass adjacent bar positions this should not affect estimates of pRF size. A slower stimulus design where each bar position is stimulated for 2-3 s, or a random instead of an ordered stimulus sequence, could probably counteract this problem but this would come at the expense of longer scanning durations. In any case, for our standard design using 1 s TR low-pass filtering clearly has no practical advantages because it does not appear to fundamentally improve map quality and skews pRF size estimates in the early visual areas.

In summary, our results show that pRF mapping with fast 1 s TR produces similar and reliable results on different scanners, even using different magnetic field strengths. It would be interesting to conduct a similar comparison between 3T and 7T scans as these are becoming increasingly commonplace.

## Acknowledgements

This work was supported by a European Research Council Starting Grant and start-up funding from the Faculty of Medical and Health Sciences at the University of Auckland. We would like to thank the staff of the Centre for Advanced Magnetic Resonance Imaging (CAMRI) for MRI support and the Centre for eResearch, University of Auckland, for computing support.

## References

Alvarez I, De Haas BA, Clark CA, Rees G, Schwarzkopf DS (2015) Comparing different stimulus configurations for population receptive field mapping in human fMRI. Front Hum Neurosci 9:96.

Amano K, Wandell BA, Dumoulin SO (2009) Visual field maps, population receptive field sizes, and visual field coverage in the human MT+ complex. J Neurophysiol 102:2704–2718.

Anderson EJ, Tibber MS, Schwarzkopf DS, Shergill SS, Fernandez-Egea E, Rees G, Dakin SC (2017) Visual Population Receptive Fields in People with Schizophrenia Have Reduced Inhibitory Surrounds. J Neurosci 37:1546–1556.

Benson NC, Jamison KW, Arcaro MJ, Vu AT, Glasser MF, Coalson TS, Van Essen DC, Yacoub E, Ugurbil K, Winawer J, Kay K (2018) The Human Connectome Project 7 Tesla retinotopy dataset: Description and population receptive field analysis. J Vis 18:23.

Brainard DH (1997) The Psychophysics Toolbox. Spatial vision 10:433–436.

Breuer FA, Blaimer M, Heidemann RM, Mueller MF, Griswold MA, Jakob PM (2005) Controlled aliasing in parallel imaging results in higher acceleration (CAIPIRINHA) for multi-slice imaging. Magn Reson Med 53:684–691.

Brown W (1910) Some Experimental Results in the Correlation of Mental Abilities1. British Journal of Psychology, 1904-1920 3:296–322.

Clavagnier S, Dumoulin SO, Hess RF (2015) Is the Cortical Deficit in Amblyopia Due to Reduced Cortical Magnification, Loss of Neural Resolution, or Neural Disorganization? J Neurosci 35:14740–14755.

Dale AM, Fischl B, Sereno MI (1999) Cortical surface-based analysis. I. Segmentation and surface reconstruction. Neuroimage 9:179–194.

de Haas B, Schwarzkopf DS (2018) Spatially selective responses to Kanizsa and occlusion stimuli in human visual cortex. Sci Rep 8:611.

de Haas B, Schwarzkopf DS, Anderson EJ, Rees G (2014) Perceptual load affects spatial tuning of neuronal populations in human early visual cortex. Curr Biol 24:R66–67.

Dumoulin SO, Hess RF, May KA, Harvey BM, Rokers B, Barendregt M (2014) Contour extracting networks in early extrastriate cortex. J Vis 14:18.

Dumoulin SO, Wandell BA (2008) Population receptive field estimates in human visual cortex. Neuroimage 39:647–660.

Edelstein WA, Glover GH, Hardy CJ, Redington RW (1986) The intrinsic signal-to-noise ratio in NMR imaging. Magn Reson Med 3:604–618.

Ekman M, Kok P, de Lange FP (2017) Time-compressed preplay of anticipated events in human primary visual cortex. Nat Commun 8:15276.

Fischl B, Sereno MI, Dale AM (1999) Cortical surface-based analysis. II: Inflation, flattening, and a surface-based coordinate system. Neuroimage 9:195–207.

Harvey BM, Dumoulin SO (2011) The Relationship between Cortical Magnification Factor and Population Receptive Field Size in Human Visual Cortex: Constancies in Cortical Architecture. J Neurosci 31:13604–13612.

Harvey BM, Dumoulin SO (2016) Visual motion transforms visual space representations similarly throughout the human visual hierarchy. Neuroimage 127:173–185.

Hoffmann MB, Kaule FR, Levin N, Masuda Y, Kumar A, Gottlob I, Horiguchi H, Dougherty RF, Stadler J, Wolynski B, Speck O, Kanowski M, Liao YJ, Wandell BA, Dumoulin SO (2012) Plasticity and stability of the visual system in human achiasma. Neuron 75:393–401.

Hughes AE, Greenwood JA, Finlayson NJ, Schwarzkopf DS (2019) Population receptive field estimates for motion-defined stimuli. Neuroimage.

Hutton C, Bork A, Josephs O, Deichmann R, Ashburner J, Turner R (2002) Image distortion correction in fMRI: A quantitative evaluation. Neuroimage 16:217–240.

Kay KN, Weiner KS, Grill-Spector K (2015) Attention reduces spatial uncertainty in human ventral temporal cortex. Curr Biol 25:595–600.

Keliris GA, Li Q, Papanikolaou A, Logothetis NK, Smirnakis SM (2019) Estimating average single-neuron visual receptive field sizes by fMRI. PNAS:201809612.

Klein BP, Fracasso A, van Dijk JA, Paffen CLE, te Pas SF, Dumoulin SO (2018) Cortical depth dependent population receptive field attraction by spatial attention in human V1. NeuroImage Available at: https://www.sciencedirect.com/science/article/pii/S1053811918303653 [Accessed May 1, 2018].

Klein BP, Harvey BM, Dumoulin SO (2014) Attraction of Position Preference by Spatial Attention throughout Human Visual Cortex. Neuron.

Kok P, Bains LJ, van Mourik T, Norris DG, de Lange FP (2016a) Selective Activation of the Deep Layers of the Human Primary Visual Cortex by Top-Down Feedback. Curr Biol 26:371–376.

Kok P, de Lange FP (2014) Shape perception simultaneously up- and downregulates neural activity in the primary visual cortex. Curr Biol 24:1531–1535.

Kok P, van Lieshout LLF, de Lange FP (2016b) Local expectation violations result in global activity gain in primary visual cortex. Sci Rep 6:37706.

Lagarias J, Reeds J, Wright M, Wright P (1998) Convergence properties of the Nelder—Mead simplex method in low dimensions. SIAM Journal of Optimization 9:112–147.

Levin N, Dumoulin SO, Winawer J, Dougherty RF, Wandell BA (2010) Cortical maps and white matter tracts following long period of visual deprivation and retinal image restoration. Neuron 65:21–31.

Moeller S, Yacoub E, Olman CA, Auerbach E, Strupp J, Harel N, Uğurbil K (2010) Multiband multislice GE-EPI at 7 tesla, with 16-fold acceleration using partial parallel imaging with application to high spatial and temporal whole-brain fMRI. Magn Reson Med 63:1144–1153.

Moutsiana C, de Haas B, Papageorgiou A, van Dijk JA, Balraj A, Greenwood JA, Schwarzkopf DS (2016) Cortical idiosyncrasies predict the perception of object size. Nat Commun 7:12110.

Moutsiana C, Soliman R, de Wit L, James-Galton M, Sereno MI, Plant GT, Schwarzkopf DS (2018) Unexplained Progressive Visual Field Loss in the Presence of Normal Retinotopic Maps. Front Psychol 9 Available at: https://www.frontiersin.org/articles/10.3389/fpsyg.2018.01722/full [Accessed October 15, 2018].

Murphy K, Bodurka J, Bandettini PA (2007) How long to scan? The relationship between fMRI temporal signal to noise ratio and necessary scan duration. Neuroimage 34:565–574.

Nelder JA, Mead R (1965) A Simplex Method for Function Minimization. The Computer Journal 7:308–313.

Schwarzkopf DS, Anderson EJ, Haas B de, White SJ, Rees G (2014) Larger Extrastriate Population Receptive Fields in Autism Spectrum Disorders. J Neurosci 34:2713–2724.

Schwarzkopf DS, Haas B de, Alvarez I (2018) SamSrf 6 - Toolbox for pRF modelling. Open Science Framework Available at: https://osf.io/2rgsm/ [Accessed August 10, 2018].

Senden M, Emmerling TC, van Hoof R, Frost MA, Goebel R (2019) Reconstructing imagined letters from early visual cortex reveals tight topographic correspondence between visual mental imagery and perception. Brain Struct Funct Available at: https://doi.org/10.1007/s00429-019-01828-6 [Accessed February 2, 2019].

Senden M, Reithler J, Gijsen S, Goebel R (2014) Evaluating Population Receptive Field Estimation Frameworks in Terms of Robustness and Reproducibility. PLOS ONE 9:e114054.

Sereno MI, Dale AM, Reppas JB, Kwong KK, Belliveau JW, Brady TJ, Rosen BR, Tootell RB (1995) Borders of multiple visual areas in humans revealed by functional magnetic resonance imaging. Science 268:889–893.

Spearman C (1904) The Proof and Measurement of Association between Two Things. The American Journal of Psychology 15:72–101.

Spearman C (1910) Correlation Calculated from Faulty Data. British Journal of Psychology, 1904-1920 3:271–295.

van Dijk JA, de Haas B, Moutsiana C, Schwarzkopf DS (2016) Intersession reliability of population receptive field estimates. Neuroimage 143:293–303.

Vo VA, Sprague TC, Serences JT (2017) Spatial Tuning Shifts Increase the Discriminability and Fidelity of Population Codes in Visual Cortex. J Neurosci 37:3386–3401.

Winawer J, Horiguchi H, Sayres RA, Amano K, Wandell BA (2010) Mapping hV4 and ventral occipital cortex: the venous eclipse. J Vis 10:1.

Winawer J, Rauschecker AM, Parvizi J, Wandell BA (2011) Population receptive fields in human visual cortex measured with subdural electrodes. Journal of Vision 11:1196.

